# Time Restricted Feeding Mitigates High-Fat-Diet Induced Sleep Disruption and Amplifies NREM Substates

**DOI:** 10.64898/2026.05.14.725282

**Authors:** Michael T. Y. Lam, Koorosh Askari, Kousha Changiz Ashtiani, Yushi Li, Nick A. Andrews, Satchidananda Panda

## Abstract

The effects of diet quality and timing on sleep quality remain poorly understood, particularly at the level of sleep microarchitecture. Traditional visual scoring captures only coarse sleep stages, overlooking the marked heterogeneity of electroencephalographic (EEG) patterns in non-rapid eye movement (NREM) sleep of mice. Here, we apply a pipeline that combines EEG feature extraction with unsupervised machine-learning-based clustering to resolve discrete NREM substates and ask how a high-fat diet (HFD) and time-restricted feeding (TRF) affect sleep microarchitectures. HFD increases sleep latency and sleep fragmentation; both abnormalities were ameliorated by active phase TRF. Clustering of 10s epochs identified two high-amplitude NREM substates sensitive to TRF: **Cluster 1**, enriched in low-delta power and peaking early in the light phase (ZT 0-6), consistent with canonical slow-wave sleep, and **Cluster 6,** characterized by elevated alpha, sigma, and beta power and peaking in the latter half of the light phase (ZT6-12). TRF increases the frequency of both NREM substates, particularly within longer uninterrupted sleep episodes during the light phases. These findings introduce an objective framework for quantifying murine sleep microarchitecture and show that aligning caloric intake with the circadian active window mitigates HFD-induced macro-level sleep disruption while selectively enhancing two physiologically distinct NREM substates.

**Significance Statement:** Time-restricted eating – targeting food intake to a defined window during the circadian active phase – confers well-established metabolic benefits, but its impact on sleep is largely underexplored. Using continuous EEG/EMG recordings, we show that an active-phase eating window mitigates high-fat-diet-induced sleep disruption in mice. We employed a novel machine-learning pipeline, further revealing that timed eating selectively increases distinct NREM substates, demonstrating that “when we eat” fine-tunes the macro- and microarchitecture of sleep. These insights lay the foundation for future translational studies and clinical trials aimed at harnessing timed eating to enhance both metabolic and sleep health.

## Introduction

Molecular circadian clocks organize and coordinate daily rhythm in activity-sleep with feeding-fasting cycle (Bass and Takahashi, 2010; Panda, 2016; Guan and Lazar, 2021). The regulation of appetite involves complex mechanisms to integrate energy storage and energy expenditure, and it must maintain a phase relationship with sleep so that hunger and foraging behavior are timed to the awake period.

Similarly, several brain regions are involved in homeostatic regulation of sleep (Chemelli et al., 1999; Lu et al., 2000; Hara et al., 2001; Adamantidis et al., 2007) and sleep regulation also integrates energy balance and food availability (Sakurai et al., 1998; Chemelli et al., 1999; Yamanaka et al., 2000; Hara et al., 2001; Dworak et al., 2010), so that sleep occurs after the animal has ingested sufficient nutrition (Jacobs and McGinty, 1971; Danguir and Nicolaidis, 1979). The energy balance state of the animal is reflected in several systemic factors, including the circulating levels of nutrients, including glucose, amino acids (Fernstrom et al., 1979), free fatty acids (Schlierf and Dorow, 1973), and ketone bodies (Iwata et al., 1991), as well as endocrine factors, including leptin (Simon et al., 1998) and ghrelin (Cummings et al., 2001; Shiiya et al., 2002). These factors exhibit diurnal rhythms that parallel the periods of feeding and fasting and their levels can affect sleep (Weikel et al., 2003; Laposky et al., 2006; Szentirmai et al., 2007). The discovery of orexin/hypocretin and its function in both appetite regulation and sleep regulation exemplifies a node in the interaction of sleep with diet (Sakurai et al., 1998; Chemelli et al., 1999; Adamantidis et al.).

Sleep and energy balance are not simple binary states. Sleep has complex macro and microarchitecture in which the latency to sleep onset, duration of sleep episodes, consolidation of sleep episodes to periods of rest are underlined by complex microarchitectures characterized by brain wave patterns such as REM and NREM sleep. Accordingly, energy state or nutrient composition can affect various aspects of sleep, but is not thoroughly understood. As changes in the quality, quantity, and timing of sleep can have pleiotropic impact on health (Belenky et al., 2003; Van Dongen et al., 2003; Scheer et al., 2009; Donga et al., 2010; Gangwisch et al., 2010; Patel et al., 2012; Roberts and Duong, 2014; Walsh et al., 2023), understanding how nutrient states affect sleep carries public health significance.

In laboratory rodent studies, specific reduction in energy intake (e.g. calorie restriction or complete fasting), changes in timing of energy availability (e.g. day-time feeding of nocturnal rodents) have shown to affect aspects of sleep-wake behaviors (Hou et al., 2019; Northeast et al., 2019; Schilperoort et al., 2019; Xu et al., 2021). Similarly, in ad libitum fed rodents, changes in macronutrient composition such as protein (Guesdon et al., 2005), or fat (Perron et al., 2015) have also shown an impact on sleep. While these studies have shed light on the impact of nutrients on sleep, they rarely address how the temporal pattern of nutrient availability affects both the macro and microarchitecture of sleep. Recent studies have shown that the temporal pattern of food intake can profoundly affect health. For example, a reduced-calorie diet given in small meals throughout the 24- hour day modestly increases the median lifespan of mice, while the same calories given in a bolus further lengthens lifespan (Acosta-Rodríguez et al., 2022). Similarly, ad libitum feeding of a high-fat diet is known to cause diet-induced obesity, while isocaloric feeding of the same high-fat diet within a <12h window (known as time-restricted feeding or TRF) prevents weight gain, metabolic diseases, and even attenuates neurodegenerative diseases in mice (Hatori et al., 2012; Chaix et al., 2014; Whittaker et al., 2023).

The TRF paradigm that partially uncouples the effects of diet on health offers an exciting opportunity to understand how diet quality and its temporal distribution affect sleep. In some human studies of Time Restricted Eating (TRE), the participants undergoing TRE also self-reported improvement in sleep quality. While there is no published study on how TRE affects sleep macro- or micro- architecture, recent report has shown that in mouse models of neurodegeneration, 6 h TRF shows modest improvement in sleep duration (Whittaker et al.). However, NREM sleep is inherently heterogeneous, with subtle variations that are not easily or reproducibly discernible using traditional scoring methods, which limits detailed characterization of how diet or any other factor affects sleep microarchitecture.

We hypothesize that diet and feeding time impact the quantity and quality of sleep, including within NREM sleep, which can be revealed through detailed analysis of EEG spectral features. Using a machine learning approach, we refined NREM staging to better characterize its dynamics and demonstrated that, under normal conditions when mice are fed a standard diet ad libitum, NREM sleep exhibits distinct electrical potentials and frequencies at different times of the day. We found that switching animals to a high-fat diet caused significant sleep disruption, including increased sleep fragmentation. Conversely, TRF reduced sleep fragmentation and increased total sleep duration, particularly during the light phase. Additionally, TRF enhanced the progression of NREM sleep during uninterrupted sleep episodes. These findings demonstrate that both diet composition and feeding schedules influence the microarchitecture of sleep. Overall, our study highlights the importance of dietary timing in modulating sleep quality and provides new insights into the interaction between metabolic health and sleep.

## Materials and Methods

### Animals

All animal experiments were carried out following the guidelines and approval of the Salk Institute IACUC. C57BL6/J male mice were purchased from Jackson Laboratory. Animals were housed in light and noise-controlled chambers set to a 12:12 hour light:dark cycle. At around 5 months of age, mice were surgically implanted with a bipotential telemetric probe for sleep recording. After surgery, they were singly housed and acclimatized in the recording chamber with autoclavable Pure-o’cell bedding (The Andersons). Unless for time-restricted feeding, animals had ad libitum access to food and water.

### Dietary Intervention and Sleep Recording

This study utilizes a pre-post experimental design to assess the effect of the timing of feeding on the quality of sleep while controlling individual differences in sleep-wake behavior. Sleep-wake states were measured using EEG/EMG on six animals across four experimental phases (Figure 1A), totaling 45 days of recording. To assess the effects of diet quality on sleep, EEG/EMG was recorded on mice fed on standard chow diet (LabDiet #5R53) (13% Fat) ad libitum for 3 days. Next, mice were fed a high-fat diet (Research Diets, Inc. D12451) containing 45% of calories from fat ad libitum (ALF-HFD). After three weeks of habituation to the HFD, sleep was recorded for three consecutive days. To test the effect of eating pattern on sleep, we subjected mice to time-restricted feeding on HFD, where the mice were provided food during their active phase (ZT13–22). Sleep was recorded after two weeks of habitation of TRF on the HFD (TRF-HFD). This feeding pattern ensures isocaloric food intake (Hatori et al.), thus isolating the effect of diet timing from nutritional quantity. To test whether the TRF effects on sleep were reversible, the mice were returned to ad libitum feeding on the HFD (post-TRF HFD), and sleep was measured after 2 weeks of habituation. Importantly, because mice handling and cage changing for food exchange occurred daily at ZT13 and ZT22 during the TRF phase, we also implemented mice handling and cage changing at ZT13 and ZT22 during the pre- and post-TRF phases of HFD feeding to control for potential investigator- or environmental- effect on sleep-wake behavior.

**Figure 1.**
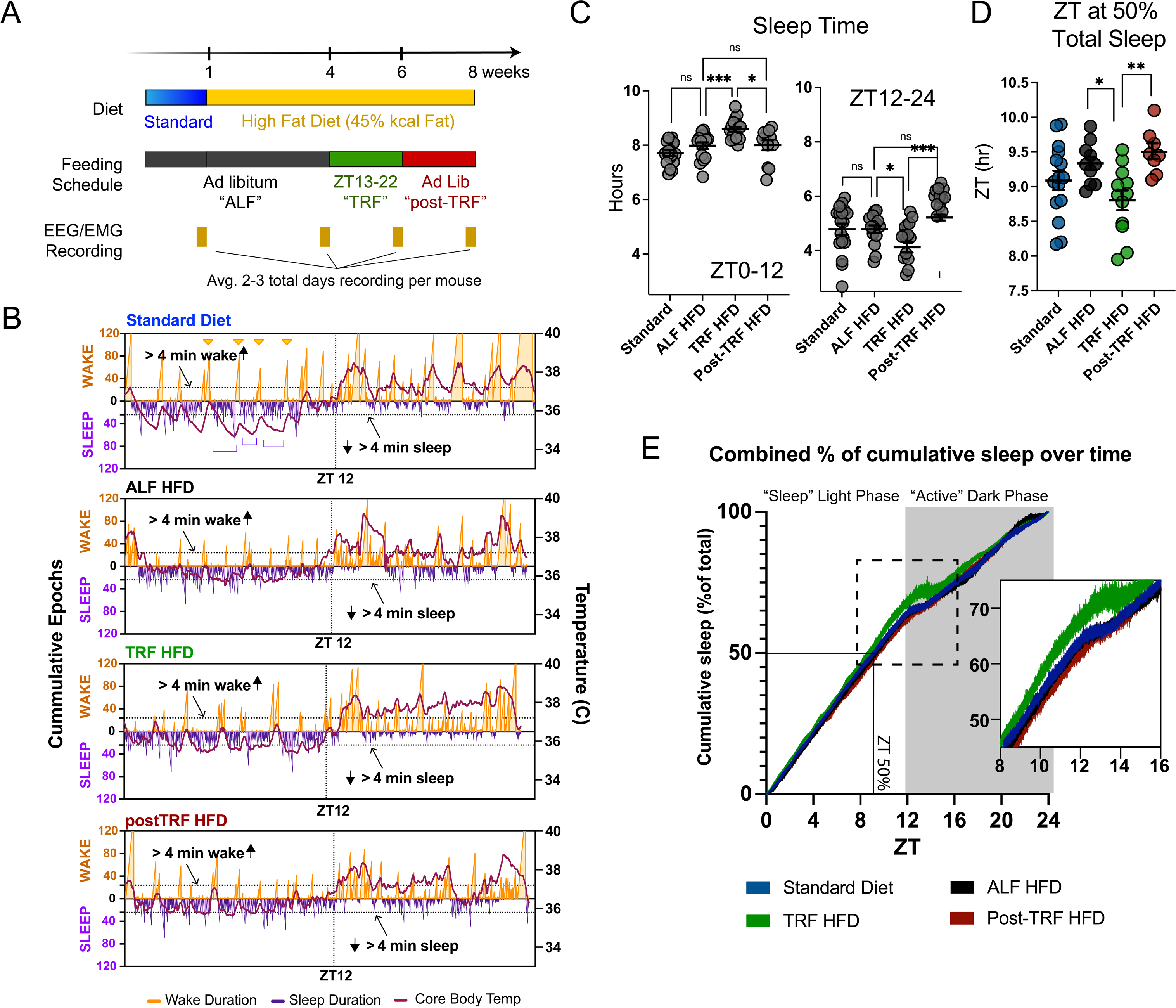
The effect of dietary intervention on sleep. A) Experimental timeline showing diet, feeding schedule, and sleep recording periods. B) Representative 24-hour sleep-wake hypnograms from each experimental phase. Sleep (violet) or wake (orange) states were assigned to every 10-second epoch starting at light exposure (ZT 0). The y-axis shows the cumulative number of uninterrupted wake (exemplified in orange triangles) or sleep episodes (exemplified in violet open brackets). The dotted line demarks 24 epochs (4 minutes). The red curve represented the core body temperature at every 10-second interval as a moving average over 5 minutes. C) Total sleep duration in the light (ZT 0-12) and dark phases (ZT 13-24). D) Zeitgeber time to 50% of total sleep. E) Cumulative sleep time across a 24-hour period. ALF: Ad libitum feeding. TRF: Time-restricted feeding. HFD: High-fat Diet. Error bars represent S.E.M. Adjusted p-value * < 0.05, ** < 0.005, *** < 0.0005, **** < 0.0001.

### Surgical Implantation of biopotential sleep recorder

EEM/EEG implantation transmitter was implanted for EEG and EMG recording in mice around 5 months of age as described previously (Ramesh et al.). Briefly, all surgical procedures were performed under sterile conditions and general anesthesia with isoflurane. First, the animals were shaved for hair removal in the ventral abdomen and dorsal neck to prepare surgical fields. The skin was sterilized using a topical application of chlorhexidine. The animal was then positioned in sternal recumbency, and a dorsal neck skin incision of 1.5-2 cm was made along the dorsal midline. A sterile bandage covered the neck incision, and the animal was turned to dorsal recumbency. A 3-4cm incision was performed through the skin and abdominal wall along the ventral midline. A telemetric transmitter (F20-EET or HD-X02, DSI, Boston, USA) was inserted into the abdominal cavity. The two pairs of biopotential leads were passed through the abdominal wall at the left mid-clavicular line using an 18- gauge needle. The wires were then exteriorized through the dorsal neck incision by subcutaneously advancing from the ventral abdomen using a trocar. The abdominal wall and skin were closed by 6-0 and 5-0 polystyrene sutures, respectively, with a simple interrupted pattern. Animals were positioned in a stereotaxic apparatus for EEG electrode implantation. The dorsal neck incision was further extended to expose the frontal area of the skull. Using a 1.0mm microdrill bit, two openings were made in the skull at the following coordinates relative to Bregma: 1.0mm anterior and 1.0mm left lateral, and 3.0mm posterior and 3.0mm right lateral. Electrodes were inserted through these openings to position them just above the cortex, underneath the skull. The electrodes were then secured by applying dental acrylic cement. The other pair of biopotentials leads was placed in contact with the muscular bundle of the right trapezius muscle to record nuchal EMG. The head and neck skin was juxtaposed with a horizontal mattress suture pattern, then closed by a simple interrupted suture pattern. Animals were given 14 days of recovery prior to sleep recording.

### Sleep Recording with EEG and EMG recording

The recording cages were placed on top of the telemetry receiver (RPC-1, DSI), which is connected to the data exchange matrix for recording (MX-2, DSI). Each mouse has at least one week of acclimatization in the cages prior to polygraphic recording. The magnetic switch of the transmitter was activated, and the physiological data were continuously acquired for at least 48 hours per condition for each animal. Dataquest ART acquisition software (include software version number) at a sampling rate of 240 Hz (F20-EET) or 300 Hz (HD-X02).

### EEG and EMG features extraction

Continuous EEG and EMG signals (0.5–25 Hz) were recorded, analyzed, and extracted in 10-second epochs using Neuroscore (DSI, software version 3.4.0). EMG amplitude was quantified using the root-mean-square of the extracted signal. For EEG, the periodogram from 0.5-25Hz was computed for total EEG power potential. We applied the Discrete Fourier Transform (DFT) to extract power in the Delta (0.5–4.0 Hz), Theta (4.0–8.0 Hz), Alpha (8.0–12.0 Hz), Sigma (12.0–16.0 Hz), and Beta (16.0–24.0 Hz) frequency ranges for each 10-second epoch, normalizing each band’s power by the total EEG power (0.5–25 Hz) to calculate the power ratio. This normalization allowed for the comparison of relative contributions of different frequency bands across varying overall signal amplitudes. To remove artifacts, epochs were excluded if they fell within the top 5% of EMG values, the extreme 2.5% of total EEG power, or the extreme 0.05% of normalized power ratios.

### Sleep Classification

Sleep-wake behavior was classified into wake, rapid eye movement (REM), and non-REM sleep using a neural network algorithm trained on manual scoring criteria derived from EMG and EEG features (Ramesh et al.). Briefly, wake epochs are characterized by high amplitude EMG and low amplitude EEG with mixed dyssynchronous frequencies. REM sleep is characterized by near absence of EMG amplitude, signifying muscle atonia, with low amplitude dyssynchronous EEG frequency. Lastly, NREM sleep is classified as low EMG amplitude with high EEG amplitude.

We used a manually scored sleep dataset from an independent mouse cohort (n = 4), totaling 16 days of recording. For the neural network algorithm, 5,000 epochs were randomly selected from each recording day. In addition, REM sleep, which only comprised about 6% of the data, was up-sampled to balance the training for wake and NREM. In total, we utilized 118,580 10-sec epochs (∼330 hours of recording), for which 80% was used for training (n = 94,839 epochs) and 20% for validation (n = 23,661 epochs).

We utilized a Deep Neural Network (DNN) to classify wake, NREM, and REM sleep based on the processed EEG and EMG features from manually scored epochs. We found satisfactory results using two hidden balanced layers (layer_specs = [7, 50, 50, 3]), with Sigmoid as the activation function and a learning rate of 0.00003. The model trained for 400 epochs, at which point the learning process converged, achieving an accuracy rate of 91.45%. Our model has 3,000 parameters, which makes the inference for the user in seconds. For output, each 10-sec epoch will have a designation of wake (W), NREM (S), and REM (P), along with the normalized EMG, EEG, and power band ratio in a csv format for downstream analysis.

### NREM Sleep Substate Analysis

NREM sleep epochs were categorized using k-means clustering based on six normalized EEG features: total power, and Delta, Theta, Alpha, Sigma, and Beta power ratios. We analyzed 166,661 10-second epochs from six male mice, setting k = 6 to generate six clusters. Cluster validation was performed by examining distinct EEG feature distributions and their temporal patterns over the 24-hour period. The clusters were labeled NREM clusters 1–6, based on their temporal phase from ZT 0. This dataset served as the discovery cohort, and new recordings were then clustered by assigning each epoch to the nearest centroid from the initial clusters, using Euclidean distance.

### Sleep Analysis

Various sleep parameters and characteristics were analyzed based on the designation of wake, REM, and NREM subclusters for each 10-second epoch. Sleep duration is defined as the total of REM and NREM clusters in defined time frames (i.e. light phase ZT 0-12, dark phase ZT 12-24, or every 2 hours). Sleep latency measures the time from light exposure (ZT0) to the first uninterrupted sleep episode lasting more than 2 minutes. Sleep arousal was defined as any sleep transition to a wake state in the subsequent epoch. Uninterrupted sleep episodes refer to sequential epochs with consecutive classifications of sleep states. The sleep-wake hypnogram depicts the sleep (violet) or wake (orange) episodes across the 24-hour time frame from ZT0, with y-axis representing the epoch number of the uninterrupted consecutive sleep or wake episodes. Temporal analyses of wake and sleep were expressed as a percentage of the recorded time within a predefined time frame. The sleep architecture at the wake-to-sleep transition was analyzed by computing the percent frequency of each NREM cluster or REM sleep for the first 15 epochs after the wake-to-sleep transition. Sleep architecture was also examined across sleep episodes of different durations and times of day (e.g., ZT 0-6, 6- 12, or 12-24). An R script for extracting uninterrupted sleep episodes based on duration or time will be made available.

## Results

### Diet conditions affect sleep duration in a time-of-day specific manner

To investigate the impact of diet quality and timing on sleep pattern, sleep was assessed from EEG and EMG recordings in male mice fed a standard chow ad libitum, HFD ad libitum or HFD-TRF (**Fig. 1A**). To visualize the effects of dietary interventions on sleep-wake pattern, actograms based on EEG and EMG recordings were generated, providing a 24-hour profile of sleep and wake pattern **(Fig. 1B)**. Sleep and wake states were assigned to each 10-second epoch, with uninterrupted sleep (purple peaks) or wake episodes (orange peaks) represented as cumulative epochs (**Fig. 1B**). The four actograms from a single mouse, representing sleep under four different diet conditions, demonstrated the overall diurnal pattern of sleep and wake behavior, with each actogram highlighting subtle yet significant characteristics (**Fig. 1B**).

Under normal chow ad libitum conditions, during the light phase (ZT 0–12) habitual sleep, multiple long clusters of sleep episodes were observed (**Fig. 1B**, purple brackets), characterized by brief interruption of wakefulness. Sustained wake episodes (usually > 4 minutes, **Fig. 1B** dotted lines) intermittently separated these larger sleep clusters (**Fig. 1B**, orange triangles). During the dark, active phase (ZT 12–24), we observed multiple large clusters of wake activity, intermittently separated by sleep clusters that typically lasted less than 4 minutes (**Fig. 1B**, dotted lines). The sequence of sleep and wake events correlated with core body temperature patterns, showing decreasing temperature during sleep and rising temperature during wake periods (**Fig. 1B**, red lines). Under ALF-HFD, there was a decrease in the long sleep clusters in the light phase. Under HFD-TRF, on the other hand, sleep was reduced in the dark, active phase. These changes observed in the actograms from a representative mouse were quantified in all mice subject to the diet conditions.

Total sleep duration was calculated based on the total number of sleep epochs during the light or dark phase (**Fig. 1C**). When fed an ad lib standard diet, the average sleep duration during the light phase was 7.7 h. which increased to 8.0 h under ad lib feeding of HFD, though this change was not statistically significant (adjusted p-value = 0.1265, mixed-effects model with repeated measures and multiple comparisons). When the mice transitioned from ad libitum HFD feeding (ALF HFD) to time-restricted feeding (TRF HFD), the average sleep duration during the light phase increased to 8.6 hours (adjusted p-value < 0.001). However, once the mice returned to ad libitum feeding of the HFD (post-TRF), their light-phase sleep duration reverted to 8.0 hours, suggesting that time-restricted feeding can increase sleep duration during the light phase in mice.

During the dark (active) phase (**Fig. 1C**, ZT 12-24), the average sleep duration was 4.8h on the standard diet and 4.9h on the ALF HFD. However, dark phase sleep duration decreased to 4.1 hours after two weeks of TRF (ALF HFD vs. TRF HFD, adjusted p-value = 0.0431) and then reverted to 5.2 hours once the mice resumed ad libitum feeding (TRF HFD vs. post-TRF HFD, adjusted p-value = 0.0003).

To assess whether diet conditions exert a time-of-day specific effect on sleep duration, the percent time spent in sleep or wake state was calculated for each 2h bin across the 24h period (**Supp Fig. 1**). Under ad lib feeding, diet quality did not significantly affect time spent in sleep during the light phase. However, HFD was associated with increased time spent sleeping at the beginning of the dark (active) phase (standard 24.4% vs. HFD 33.0 % ZT 12-14, adjusted p-value 0.0205) and reduced time spent in sleep at the end of the dark phase (standard 33.0% vs. HFD 15.8%, ZT 22-24, adjusted p-value < 0.0001). TRF affected sleep during both light and dark phases. Compared to the mice on ALF HFD, the TRF HFD feeding condition increases time spent in sleep during the early light phase (ZT 0-2, ALF HFD 62.0%, TRF HFD 74.9% adjusted p-value 0.002). We tabulate the hours it takes to reach 50% of all sleep within a 24 hour period. For ALF standard, half of the total 24-hour sleep was reached at ZT 9.1 (**Fig. 1D**). For ALF-HFD, ZT 9.3. Mice on TRF-HFD reached 50% of total sleep within 24hr at 8.8 hr compared to ALF-HFD (ZT 9.3, adjusted p value = 0.0389) or post-TRF HFD (ZT 9.5, adjusted p-value 0.0076, **Fig. 1D**), suggesting mice consolidate more sleep during the light phase in TRF condition (**Fig. 1E**). Conversely, TRF significantly decreased time spent in sleep during the active phase at ZT 18-20 (ad lib 56.7%, TRF 40.3% adjusted p-value < 0.0001) and 20-22 (ad lib 52.4%, TRF 38.0% adjusted p-value < 0.0001) (Supp Fig. 1). This reduction in sleep during the active phase was accompanied by a corresponding increase in core body temperature, indicating heightened wakefulness (**Fig. 1B**).

### Effect of diet conditions on sleep quality

When allowed to sleep, the time it takes to fall asleep and the quantity of sleep arousal constitute sleep quality. The average time for sleep onset, defined as the time from lights on at ZT 0 to the first episode of uninterrupted sleep ≥2 minutes (**Fig. 2A**), did not change significantly when mice were fed a standard diet or ALF HFD (17.9 min standard diet, 23.7 min HFD, adjusted p-value = 0.79). Under TRF, sleep onset significantly decreased (23.7 min ALF HFD to 10.2 min TRF, adjusted p-value = 0.0168, **Fig. 2A**). These findings suggest TRF shortens sleep onset time.

**Figure 2.**
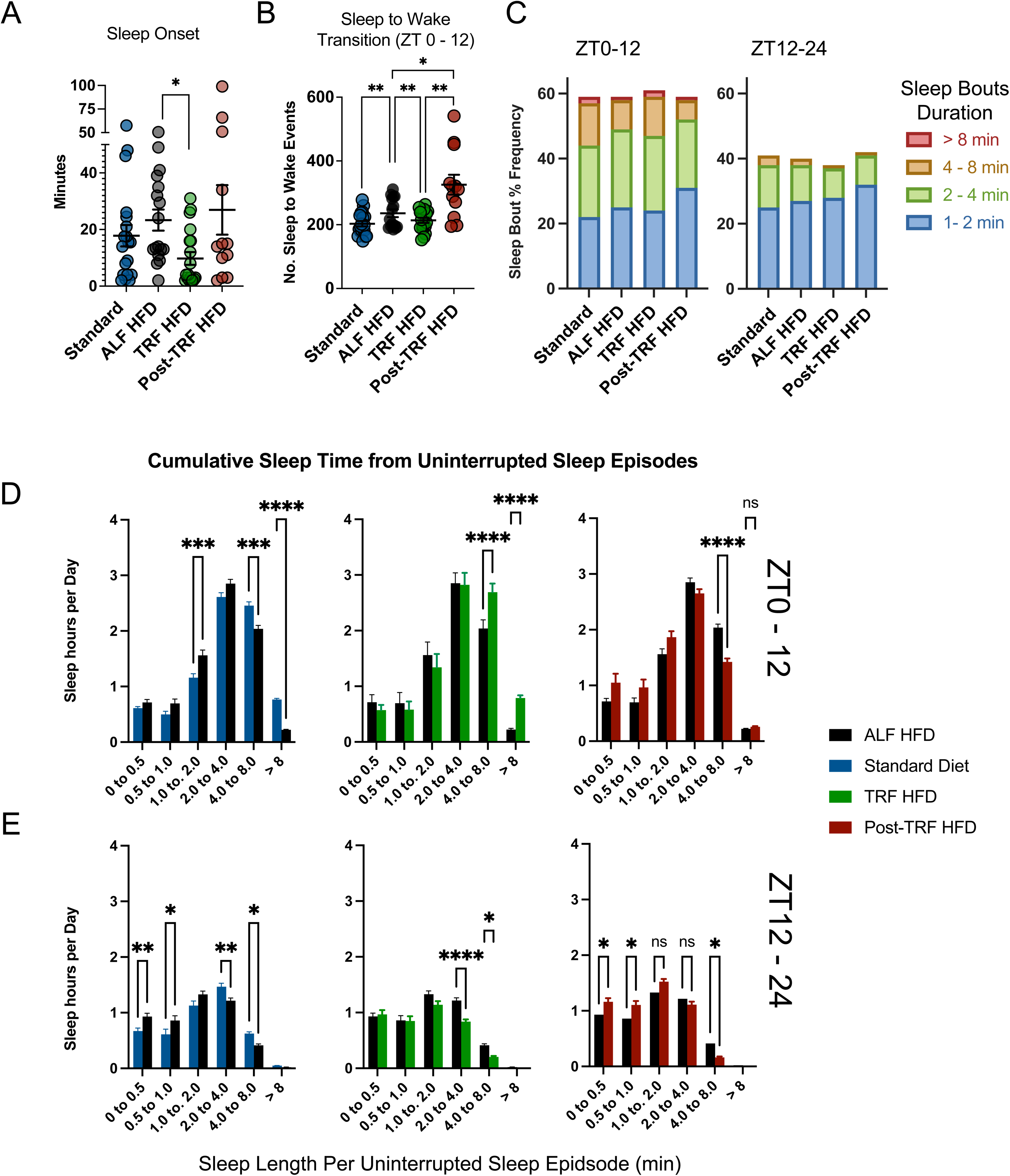
Dietary Effects on Sleep Quality. **A)** Sleep onset, defined as the time (in minutes) from light exposure (ZT 0) to the first uninterrupted sleep episode lasting at least 2 minutes. **B)** Number of sleep-to-wake transitions (sleep arousals) between ZT 0–12. **C)** Percentage distribution of sleep episodes based on uninterrupted sleep duration. **D-E)** Cumulative sleep time (in hours) from sleep episodes of varying uninterrupted durations during the **D)** light phase (ZT 0–12) and **E)** dark phase (ZT 12–24). Adjusted p- value * < 0.05, ** < 0.005, *** < 0.0005, **** < 0.0001.

We next evaluate sleep fragmentation by examining the frequency of sleep-to-wake transitions (**Fig. 2B**). During the light phase, animals on ALF HFD exhibited an increased number of sleep-to-wake transitions (average 203 standard diet vs 250 ALF HFD adjusted p-value = 0.0002). However, TRF reduced the sleep-to-wake transition frequency to 202 (p-value = 0.002, ALF HFD vs. TRF HFD). Consistently, the frequency of sleep-to-wake transitions increased again after the animals returned to the ad libitum feeding schedule, suggesting that the effect is specific to time-restricted feeding. Overall, these findings indicate that a dietary change from standard chow to a high-fat diet increases sleep arousal or sleep fragmentation, an effect that is mitigated by time-restricted feeding.

Another sleep quality surrogate is the length of uninterrupted sleep episodes or sleep bouts (**Fig. 2C**). Sleep episodes were binned into four groups: short (1–2 min), medium (2–4 min), long (4–8 min), and very long (> 8 min). Under ad lib feeding conditions, HFD increased the proportion of short and medium length sleep bouts while reducing long and very long sleep bouts (standard diet; 49, 50, 29, 5 bouts vs. HFD; 64, 60, 23, 2 short, medium, long and very long bouts respectively, **Fig. 2C and Supp Fig. 2**). Conversely, TRF HFD reversed this trend, and the mice regained the long and very long sleep bouts (**Fig. 2C, Supp Fig. 2**). In accordance with the change in distribution of sleep bout lengths, the fraction of total sleep attributed to various sleep bout lengths also changed (**Fig. 2D-E**). During the light (sleep) phase, ALF-HFD mice had significantly more sleep from short bouts than the same mice fed ad lib a standard diet. TRF-HFD reversed this trend, increasing the total sleep time in more prolonged sleep bouts (4-8 minutes and > 8 minutes, **Fig. 2D**).

During the dark (active) phase, HFD reduced the length of time spent in long and very long sleep bouts, and the mice also spent more time under very short sleep bouts lasting <1min (**Fig. 2E**). TRF further reduced the duration of time spent in long and very long sleep bouts.

### Effects of Diet Condition on REM and NREM Sleep

Next, we examined the effects of diet and feeding schedule on Rapid Eye Movement (REM) and Non-REM (NREM) sleep. When fed ad lib, the diet quality did not affect overall NREM or REM sleep during either the light or dark phase (**Fig. 3A & B**, standard vs. HFD, light phase NREM sleep 6.9 vs 7.0 hours; dark phase NREM sleep 4.3 vs. 4.4 hours; light phase REM 0.83 vs 0.85 hours; and dark phase REM sleep 0.44 and 0.48 hours; adjusted p-values > 0.05). However, time-restricted feeding of HFD led to a modest but significant increase in NREM and REM sleep during the light phase by 22.8 and 2 minutes, respectively (**Fig. 3A & B**, adjusted p-values 0.008 and 0.008). In contrast, during the active phase, TRF reduced REM sleep by 10 minutes (**Fig. 3B**, adjusted p-value = 0.0076). We also evaluated the percentage of NREM and REM sleep in 2 hr bins. A significant decrease in NREM sleep was observed between ZT18–22 under TRF-HFD compared to ad lib feeding of HFD (adjusted p-values = 0.005 and 0.0114, respectively), with a trend toward increased NREM sleep during the early sleep phase (ZT0–2) (adjusted p-value = 0.065) (**Fig. 3C, middle**). Overall, the impact of TRF on NREM and REM sleep was modest (**Fig. 3D**).

**Figure 3.**
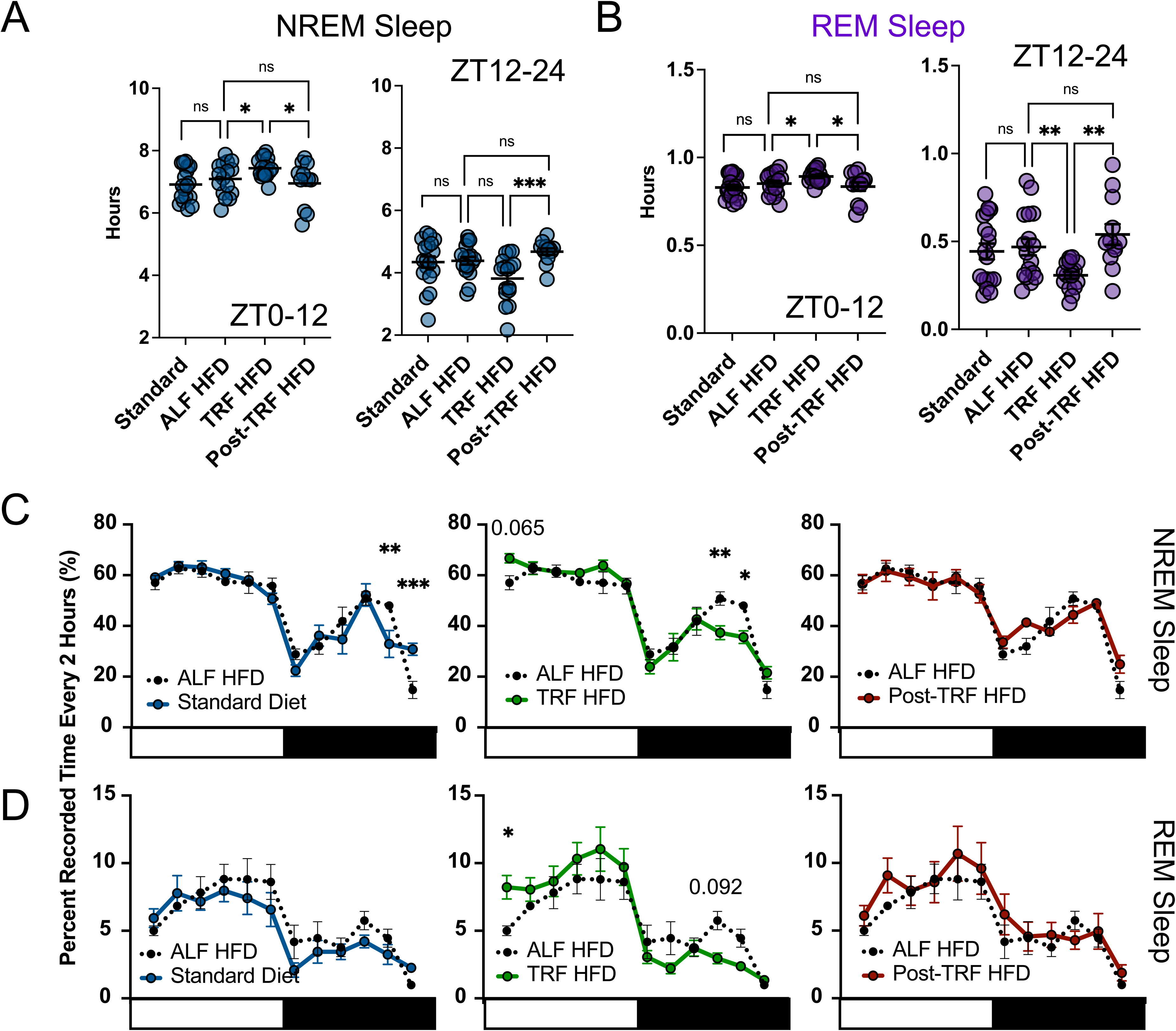
Dietary Effects on NREM and REM Sleep. Cumulative **A)** NREM and B) REM sleep duration in ZT 0–12 and ZT 12–24 across the four phases of the dietary interventions. **C-D)** Temporal distribution of **C)** NREM and **D)** REM sleep every two hours across the 24-hour period. Standard is shown in blue, ALF HFD in dotted black, TRF-HFD in green, and Post-TRF-HFD in red. Error bars represent S.E.M. Adjusted p-value * < 0.05, ** < 0.005, *** < 0.0005, **** < 0.0001.

### Machine learning approach for NREM sleep classification

We hypothesized that the impact of diet and TRF on NREM sleep structure might be obscured by the inherent heterogeneity within NREM sleep, which, by definition, includes any sleep episode that is not REM. During manual scoring with EEG waveform visualization, we frequently observed distinct patterns within NREM sleep, including high-amplitude slow waves (∼4.0 Hz), lower amplitude waves with higher frequencies (>12 Hz), and high-amplitude waves with mixed frequencies (**Fig. 4A**). Discerning these subtle waveform patterns through visual inspection is challenging, prone to subjective bias, and often lacks reproducibility. This complexity suggests that treating NREM sleep as a single entity may oversimplify its structure and lead to the oversight of important biological effects. To address this gap and ensure objective and reproducible analysis of NREM sleep heterogeneity, we employed unsupervised machine learning approaches to identify distinct subgroups of NREM sleep based on EEG characteristics (**Fig. 4B**). This method allows us to more accurately capture how these dietary interventions might differentially affect the various subtypes of NREM sleep.

**Figure 4.**
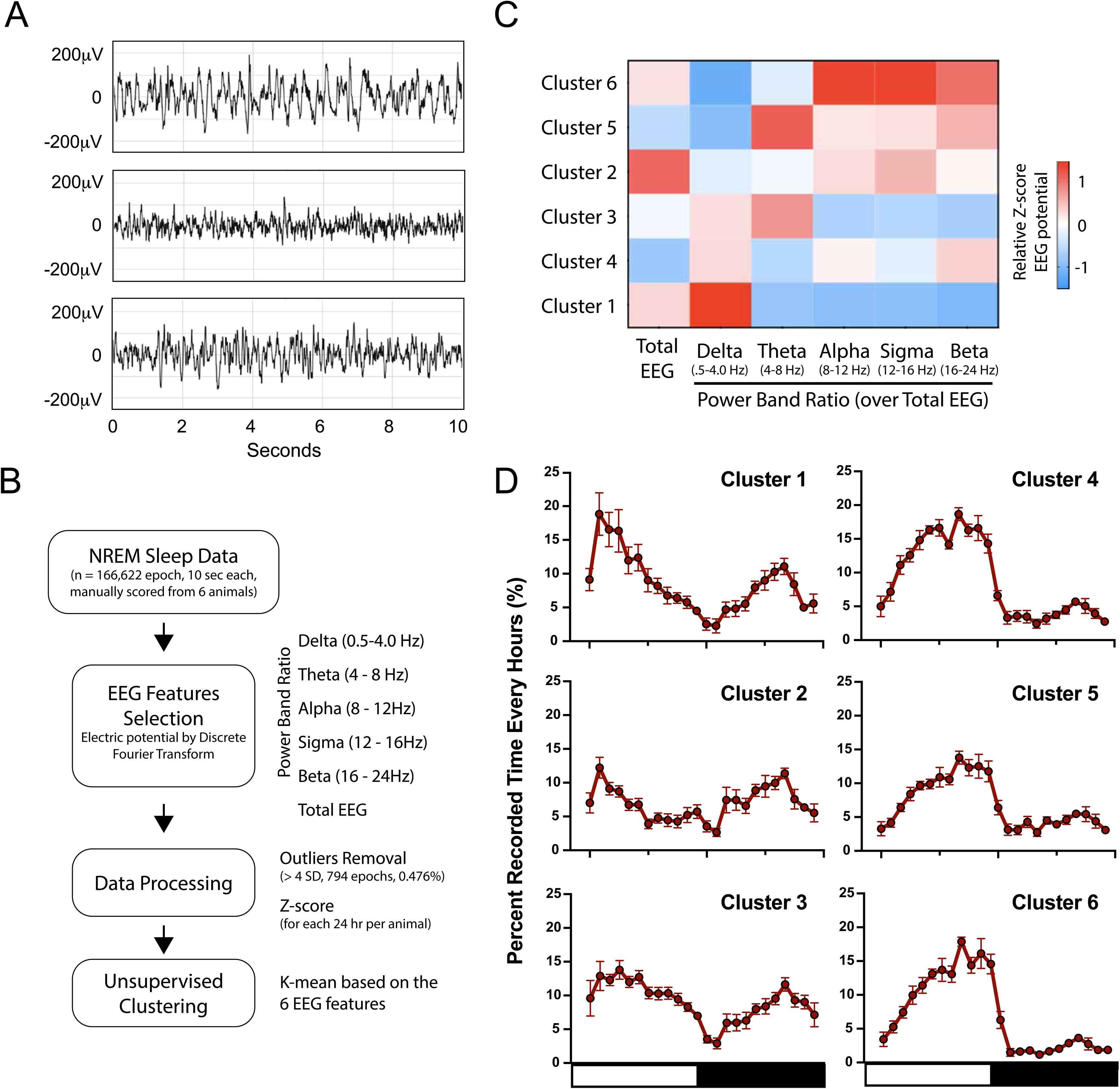
Identification of NREM Subgroups Using Unsupervised Clustering. **A)** Representative 10-second EEG traces from different NREM sleep epochs showing distinct amplitude and frequency characteristics. **B)** Workflow illustrating the steps for clustering NREM sleep, including data selection, feature extraction, preprocessing, and unsupervised clustering. **C)** Heatmap displaying relative EEG power and the Delta, Theta, Alpha, Sigma, and Beta ratios across six NREM clusters, with red indicating higher values and blue lower values. **D)** Temporal distribution of NREM Clusters 1–6 across 24 hours. The white bar indicates the light phase (ZT 0–12), and the black bar indicates the dark phase (ZT 12–24). Error bars represent S.E.M.

To ensure reproducibility in identifying NREM subgroups through unsupervised machine learning, sleep recordings from an independent cohort of mice (n = 6) were used. For each 10-second epoch, the total EEG power across the frequency range of 0.5–25 Hz was extracted using Discrete Fourier Transform (DFT). This total EEG power was normalized to the average power across all epochs within the 24-hour recording period, reflecting the relative EEG power throughout the day. Additionally, using DFT, electrical potential from specific frequency bands—delta (0.5–4.0 Hz), theta (4–8 Hz), alpha (8–12 Hz), sigma (12–16 Hz), and beta (16–24 Hz)— was extracted and normalized each to the total EEG power as a Power Ratio. This method provides interpretable results, as sleep is traditionally characterized by waveforms within the delta and theta frequency ranges. Consequently, each NREM epoch can be described by six numerical features: its overall EEG activity and the relative EEG power potential across the five frequency bands (**Fig. 4B**).

We employed unsupervised machine learning to identify NREM sleep subgroups based on six EEG features, using K-means clustering on 166,622 10-second NREM sleep epochs (6 mice, ∼460 hours of recordings). To determine the optimal number of clusters, we calculated silhouette scores, which indicated an ideal cluster number between 6 and 8 (silhouette score 0.1732). We then characterized the distribution of these NREM sleep clusters across the 24h period, reasoning that clusters with biological significance should exhibit non-random 24h rhythms. This approach identified six distinct NREM sleep clusters, each exhibiting unique amplitude patterns across the five frequency bands (**Fig. 4C**), each exhibiting specific temporal abundance across the 24h period (**Fig. 4D**). For example, Cluster 1, characterized by a high power ratio in the low-frequency delta range, is predominantly observed during the early sleep phase, aligning with slow-wave sleep. Clusters 2 and 3 also peak during the early sleep phase, while Clusters 4, 5, and 6 peak later in the sleep phase. Notably, Cluster 6, which ranks as the second highest in total EEG potential, shows a power ratio predominantly in the higher frequency bands – Alpha, Sigma, and Beta (**Fig. 4C**).

Crucially, we identified similar NREM clusters with consistent temporal distribution in the Diet and TRF mouse cohort (**Supp Fig. 3**), underscoring the reproducibility of our approach. This robust scoring scheme enables the objective quantification of different NREM subgroups, allowing for a detailed sleep examination at a finer resolution.

### Sleep Architecture at Wake to Sleep Transition

To explore the potential biological implications of the NREM clusters, we examined sleep at a finer temporal resolution by focusing on wake-to-sleep transitions. We analyzed the first 15 epochs (2.5 minutes) following a wake-to-sleep transition in uninterrupted sleep bouts lasting 2.5 to 4 minutes (**Fig. 5A**). The frequencies of Clusters 1, 2, and 6 increase early in the wake-to-sleep transition, reflecting their rising frequency in the initial phases of sleep as these clusters become more prominent as the uninterrupted sleep progresses (**Fig. 5B**). Conversely, Clusters 4 and 5 decrease early in the transition, while Cluster 3 shows a more gradual decrease (**Fig. 5C**). Clusters 2, 6, and 1 rank highest in total EEG potential, in that order, while Clusters 4 and 5 rank lowest (**Fig. 5D**), suggesting an increase in EEG power as the sleep episode progresses. As the sleep episode moves toward wakefulness, REM sleep gradually increases, reflecting its role in the latter stages of the sleep cycle.

**Figure 5.**
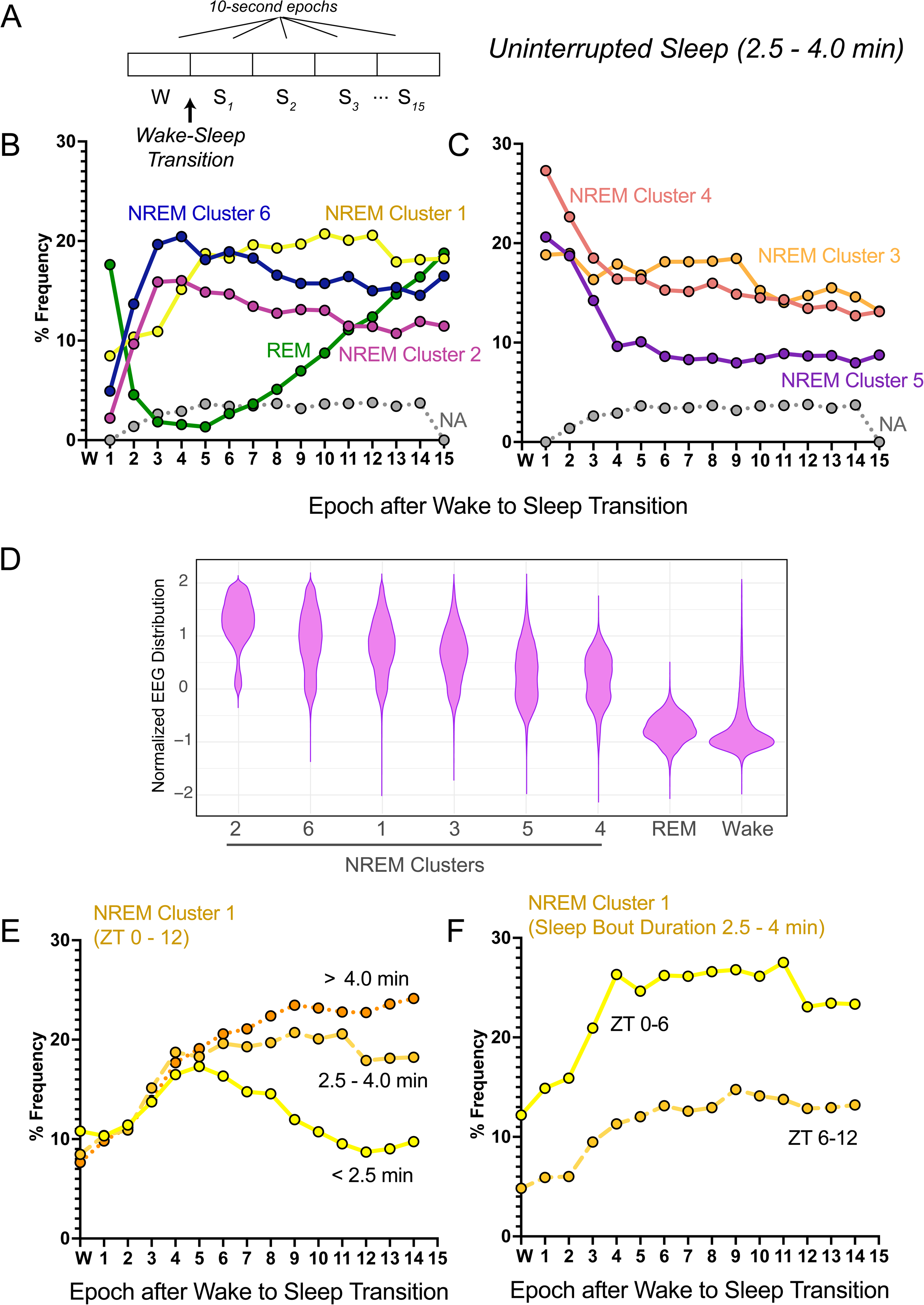
Sleep Architecture at the Wake-to-Sleep Transition. A) Graphical representation of a wake-to-sleep transition, where W represents wake, Sn indicates sleep, and n refers to the epoch number after the transition. Sleep architecture analysis focuses on episodes lasting 2.5 to 4 minutes. B-C) Relative frequency of sleep states for the first 15 epochs post-transition during ZT 0-12. B) NREM Clusters 1, 6, 3, and REM increase in relative abundance, while C) NREM Clusters 2, 4, and 5 decrease as sleep progresses. NA represents missing data from epochs filtered out during the preselection process. D) Violin plot showing normalized total EEG power, where NREM Clusters 2, 6, and 1 exhibit the highest values. E) Relative frequency of NREM Cluster 1, characterized by high EEG amplitude in Delta frequency, during the wake-to-sleep transition for uninterrupted sleep episodes of <2.5 minutes, 2.5–4 minutes, and >4 minutes during the light phase (ZT 0-12). F) Comparison of NREM Cluster 1 frequency during uninterrupted sleep episodes (2.5–4 minutes) between the early (ZT 0–6) and late (ZT 6–12) sleep phases.

We then focused on Cluster 1, characterized by EEG patterns with the highest prevalence of delta low-frequency waveforms, particularly early in the sleep phase (ZT 1), resembling slow-wave sleep. We analyzed Cluster 1’s behavior across sleep bouts of varying durations and phases of the sleep cycle. In shorter sleep bouts (1–2.5 minutes), Cluster 1 peaks five epochs after the wake-to-sleep transition, coinciding with the epoch when some bouts transition back to wakefulness (**Fig. 5E**). In longer sleep bouts (>4 minutes), the frequency of Cluster 1 continues to increase by epoch 15 and is higher than its frequency in sleep bouts lasting 2.5 to 4 minutes. Noting that Cluster 1 peaks at ZT 1 across the 24 hours (**Fig. 4D**), we further investigated whether the pattern of Cluster 1 at the wake-to-sleep transition varies at different times within the light phase. We compared the frequency of Cluster 1 at the wake-to-sleep transition in sleep bouts lasting 2.5 to 4 minutes between ZT 0–6 and ZT 6–12. The frequency of Cluster 1 was higher during the early phase of sleep (ZT 0–6) (**Fig. 5F**). This approach reveals that Cluster 1 accumulates as the uninterrupted sleep episode progresses and underscores the heterogeneity of sleep architecture across different phases of the sleep cycle.

### Time-of-the-day specific Sleep Episode Composition

Next, we investigated the composition of sleep at the wake-to-sleep transition at different times of the day, based on our observation that the NREM clusters peak at different points during the 24-hour period. In early light phase (ZT 0 – 6), among the NREM clusters that increase as the uninterrupted sleep episode progresses, Cluster 1 peaks at the 5th epoch and is the predominant NREM cluster (**Fig. 6A**). In the later light phase (ZT 6–12), Cluster 6 peaks at the 3rd epoch and becomes the dominant NREM subgroup as sleep progresses during this time frame (**Fig. 6B**). Both Clusters 1 and 6 have high total EEG potential (**Fig. 5D, Supp Fig. 3**), with Cluster 1 showing a high Delta power ratio and Cluster 6 exhibiting greater power in the higher frequency bands (**Fig. 4C, Supp Fig. 3**). During the dark, active phase (ZT 12–24), we found that Cluster 2 is more abundant than Cluster 1 or 6 during the ‘*siesta*’ period (**Fig. 6C**).

**Figure 6.**
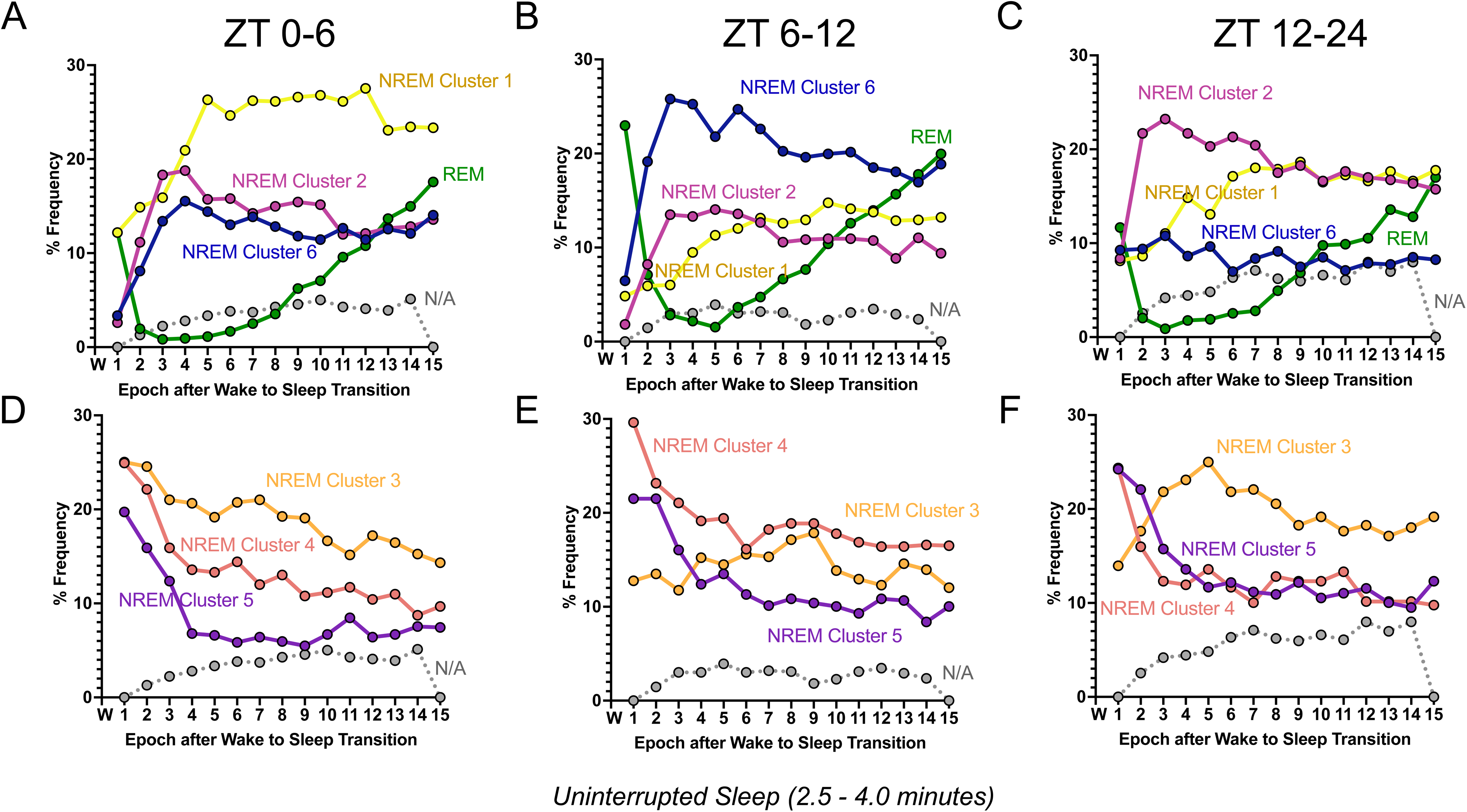
Time-of-Day Differences in Sleep Architecture at Wake-to-Sleep Transitions. The relative abundance of sleep states during the first 15 epochs following wake-to-sleep transitions in uninterrupted sleep episodes (2.5–4.0 minutes). A-C show the sleep states that increase in abundance as sleep progresses, while D-F show NREM clusters that generally decrease. A & D represent early light phase (ZT 0–6), B & E represent late light phase (ZT 6–12), and C & F represent dark phase sleep (ZT 12–24). The relative abundance of sleep clusters differs across these time intervals.

The different times of day also influence the NREM clusters that descend as sleep episode progresses. Cluster 3 is more abundant than Clusters 4 or 5 in the early sleep phase (ZT 0–6, **Fig. 6D**), while Cluster 4 becomes more abundant than Clusters 3 or 5 in the latter part of the light phase (ZT 6–12, **Fig. 6E**). During the *siesta* period in the dark phase, NREM Cluster 3 increases in frequency early in the wake-to-sleep transition, contrasting with its behavior during sleep episodes in the light phase (ZT 0–6 and ZT 6– 12) (**Fig. 6F**). These analyses underscore that sleep composition varies significantly depending on the time of day, highlighting the dynamic nature of NREM clusters and their potential role in sleep architecture.

### Diet conditions affect NREM subclusters

Next, we evaluated the effects of dietary changes and the timing of feeding on the NREM subclusters. Given that the NREM clusters are distributed differently throughout the day, we analyzed changes in the abundance of these clusters during the early sleep phase (ZT 0–6), late sleep phase (ZT 6–12), and the active phase (ZT 12–24). We first examined the effects of transitioning mice from a standard chow diet to an HFD on the NREM clusters. No NREM clusters showed sustained changes in abundance when comparing standard chow to ALF HFD or post-TRF HFD conditions (**Fig. 7**).

**Figure 7.**
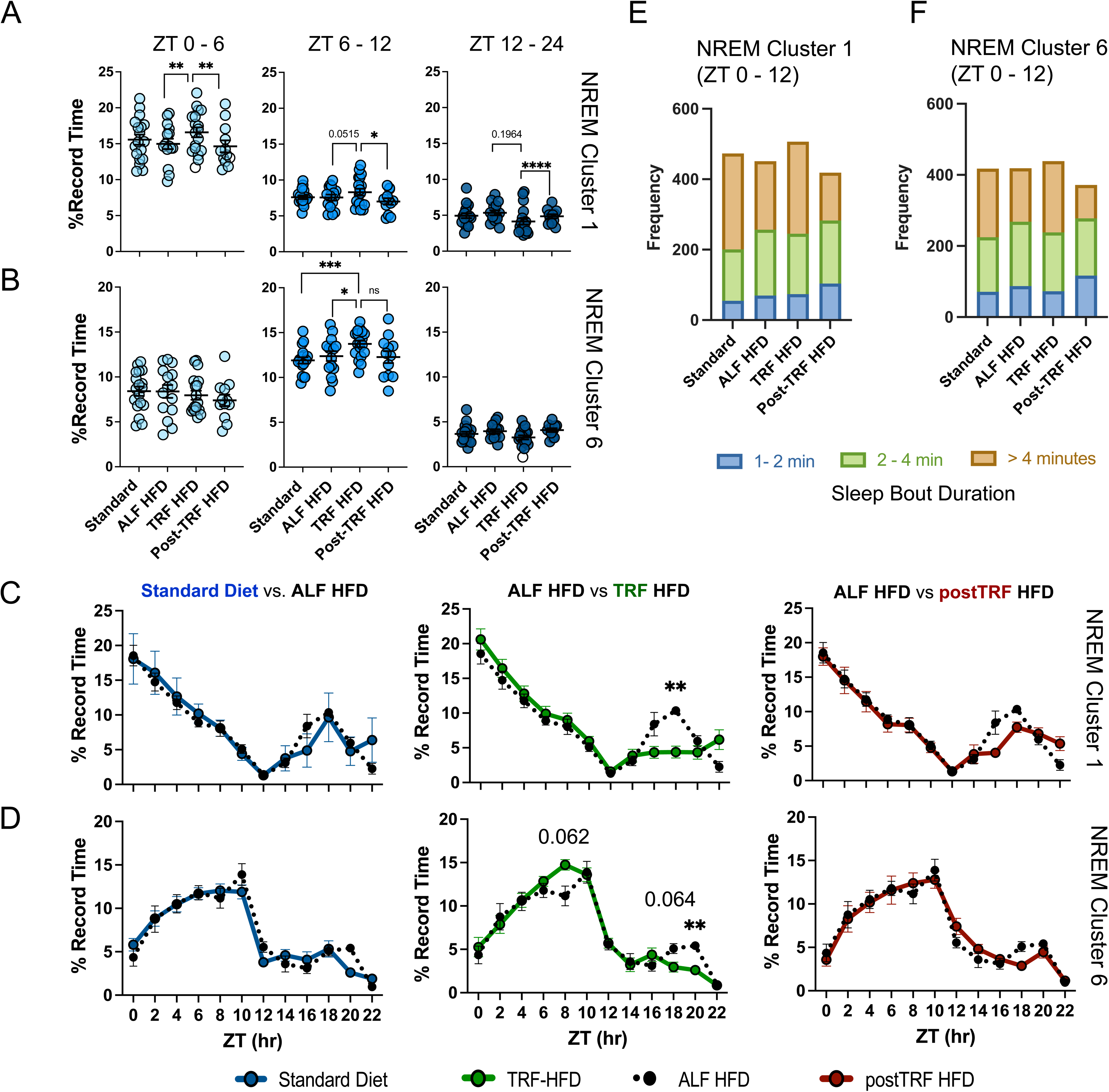
Effect of Dietary Intervention on NREM Clusters 1 and 6. A-B) Abundance of A) NREM Cluster 1 and B) NREM Cluster 6 across experimental phases for each animal in a pair-wise fashion. C-D) Temporal distribution of C) NREM Cluster 1 and D) NREM Cluster 6 over 24 hours. Standard chow is shown in blue, ALF HFD in dotted black, TRF-HFD in green, and Post- TRF-HFD in red. Error bars represent S.E.M. E-F) Average frequencies of E) NREM Cluster 1 and F) NREM Cluster 6 for sleep episodes of 1–2 minutes (blue), 2–4 minutes (green), and >4 minutes (brown). Transition from standard chow to HFD reduced NREM Cluster 1 and 6 during uninterrupted sleep >4 minutes, while TRF increased longer sleep episodes. Adjusted p-value * < 0.05, ** < 0.005, *** < 0.0005, **** < 0.0001.

We then examine the effect of TRF on NREM sleep clusters. Among all the NREM clusters, TRF specifically affected Clusters 1 and 6, which are the ascending NREM sleep clusters as a sleep episode progresses. Cluster 1 resembles slow-wave sleep and is the dominant ascending NREM cluster in an uninterrupted sleep episode with high EEG potential in the Delta band during the early sleep phase (ZT 0–6; **Fig. 4 and 6**). Cluster 6, on the other hand, is the dominant ascending NREM cluster in an uninterrupted sleep episode with high EEG potential in the higher frequency bands during the late sleep phase (ZT 6–12; **Fig. 4 and 6**). When the mice were on an HFD, the TRF paradigm significantly increased the abundance of Cluster 1 during the early sleep phase (**Fig. 7A**, ZT 0–6; adjusted p-value = 0.0036). However, the effect of TRF on Cluster 1 was less pronounced during the late sleep phase (**Fig. 7A**, ZT 6–12). In contrast, TRF’s effect on Cluster 6 was notable during the late sleep phase (**Fig. 7B**, ZT 6–12) but not during the early sleep phase (**Fig. 7B**, ZT 0–6).

We further examined the frequency of Clusters 1 and 6 at 2-hour bins across the 24-hour period. A significant decrease in Cluster 1 was observed at ZT 18 during the siesta period when the mice were on TRF HFD (**Fig. 7C**, ALF-HFD vs TRF HFD, adjusted p-value = 0.008). Similarly, TRF decreased Cluster 6 during the siesta, consistent with the overall reduction in sleep time during the active phase (**Fig. 7D**, preTRF vs TRF HFD).

We then analyzed the abundance of Clusters 1 and 6 in sleep bouts of different lengths (**Fig. 7E & F**). In this analysis, we observed a decrease in Clusters 1 and 6 in sleep bouts lasting longer than 4 minutes when the diet was changed from standard chow to HFD during ZT 0–12, despite the average total frequency of Cluster 1 not being significantly different between the two groups (**Fig. 7A, C, and E**). TRF increased the abundance of Clusters 1 and 6 in sleep bouts longer than 4 minutes, but this effect reversed in the post-TRF HFD group (**Fig. 7E & F**).

Together, our findings demonstrate that HFD and TRF have specific and distinct effects on Clusters 1 and 6, highlighting the nuanced impact of diet and feeding timing on sleep architecture.

## Discussion

Here, we demonstrated the quantitative effects of dietary interventions on sleep quality and architecture. Switching the animals from a 13% fat diet to a 45% high-fat diet worsens sleep quality by increasing sleep fragmentation, consistent with previous studies (Panagiotou et al., 2018; Panagiotou and Deboer, 2019). However, within our experimental parameters, we did not observe an increase in total sleep time due to the high-fat diet reported in earlier studies (Jenkins et al., 2006), suggesting that HFD does not necessarily alter total sleep duration.

We found that two weeks of TRF reverses HFD-induced sleep fragmentation. This effect is consistent with our previous metabolic studies, where aligning feeding to the active phase reversed HFD-driven metabolic impairments, even when caloric intake was equivalent between groups (Hatori et al., 2012; Chaix et al., 2014; Chaix et al., 2019; Deota et al., 2023). Similarly, TRF increased total sleep time during the light phase while reducing sleep during the active phase. Notably, the timing of food intake also reduces sleep fragmentation in mouse models of Alzheimer’s and Huntington’s disease (Whittaker et al., 2023; Chiem et al., 2024), suggesting that timed feeding not only promotes consolidated sleep but does so when sleep is aligned with the circadian rest phase.

In this manuscript, we quantified subgroups of NREM sleep by leveraging EEG features and a machine learning approach to objectively characterize the heterogeneity of NREM sleep. Specifically, we classified NREM sleep based on EEG potential amplitude and the relative power across frequency bands (Delta, Theta, Sigma, Alpha, and Beta) using unsupervised K-means clustering. This method captures subtle EEG dynamics and identifies six distinct NREM subgroups, each showing varying temporal abundance across the 24-hour period, suggesting potential biological roles for each subgroup. Our analysis highlights NREM Cluster 1, which aligns with slow-wave activity based on power spectral analysis. Cluster 1 is characterized by high EEG potential amplitude and a strong Delta band component (0.5–4.0 Hz). Its occurrence peaks at the beginning of the sleep phase, resembling slow-wave sleep (SWS) described in humans (Berry et al., 2018) and rodents (Wei et al., 2019).

A key strength of our analysis is that it examines sleep microarchitecture in a contiguous fashion—tracking the prevalence of different NREM subgroups continuously across uninterrupted sleep episodes. This represents a novel approach to characterizing sleep structure beyond conventional epoch-based scoring. Following the wake-to-sleep transition, we observed that NREM Cluster 1 increases frequency as the uninterrupted sleep episode progresses. Notably, the abundance of Cluster 1 at the wake-to-sleep transition varies depending on the time of day and the duration of the sleep episode. During the early sleep phase (ZT 0–6), Cluster 1 is the primary accumulating NREM subgroup. In the late sleep phase (ZT 6–12), NREM Cluster 6—characterized by elevated EEG potential amplitude in higher frequency bands (Sigma, Alpha, and Beta)—becomes more prominent. During naps in the active phase (ZT 13–24), NREM Clusters 2 and 3 are more prevalent.

To our knowledge, our quantitative approach to characterizing the microarchitecture of continuous sleep in rodents is unique and complements existing methods for understanding how different behavioral contexts influence sleep. From this perspective, we found that both diet and TRF specifically impact NREM Clusters 1 and 6—subgroups of NREM sleep whose frequency increases as an uninterrupted sleep episode progresses. TRF increases the frequency of NREM Cluster 1 during the early sleep phase (ZT 0–6) and Cluster 6 during the late sleep phase (ZT 6–12). Notably, these increases are most pronounced in uninterrupted sleep episodes longer than 4 minutes, suggesting that Clusters 1 and 6 may contribute to consolidated sleep.

We also observed that dietary changes, particularly transitioning from standard chow to HFD, alter Clusters 1 and 6 depending on the duration of uninterrupted sleep episodes. Under HFD, the frequency of Clusters 1 and 6 decreases in sleep episodes longer than 4 minutes, even though the total frequency of Cluster 1 or 6 across all sleep episode durations does not significantly change. These findings suggest that examining sleep episodes of different lengths can uncover subtle shifts in sleep architecture and reveal how diet and feeding schedules modulate specific aspects of sleep.

Our approach complements existing sleep analytical tools. The traditional power spectral analysis examines average electrical potential of across frequency bands over broader time frames (such as each hour or each sleep cycle), our approach complements this by analyzing the characteristics of NREM clusters at a finer resolution for each 10- second epoch. Additionally, our approach builds on prior work in automated sleep-wake classification in larger rodents (Kohtoh et al., 2008; Stephenson et al., 2009; Miladinović et al., 2019; Wei et al., 2019). It would be interesting to assess how our NREM subgroups align with these earlier automated classifications (e.g. transitional sleep), particularly since rat studies benefit from simultaneous multi-site EEG recordings (e.g., hippocampal and occipital channels), which may provide additional resolution.

We also acknowledge that while features such as sleep spindles and K-complexes typically fall within the Sigma frequency band, our current approach was not optimized to capture these discrete events. Detecting such features may require analyzing sleep at a finer temporal resolution (e.g. < 2 second per epoch) or with different approach (e.g. wavelet analysis). Finally, it will be valuable to explore how different biological perturbations affect other NREM subgroups to better understand their distinct biological roles.

In summary, using machine learning to analyze distinct NREM EEG and EMG characteristics, we developed a method to objectively evaluate the effects of diet and feeding timing on sleep behavior in a mouse model. Given the heterogeneity observed in human studies, and the association of specific EEG features with sleep-related disorders (Kang et al., 2020; G. Horváth et al., 2022; Kang et al., 2022), accurate tools in animal models that can focus on subtle but reproducible EEG patterns are crucial for understanding how diet and metabolism influence sleep behavior.

## Acknowledgments

The authors acknowledge the technical and logistical help from Hiep Le and Arshia Farajnejad. The study was supported by the Academic Sleep Pulmonary Integrated Research/Clinical Fellowship through the American Thoracic Society, Veterans Affairs Biomedical and Laboratory Research and Development Career Development Award (1 IK2 BX0059089), and by NIH (5T32HL134632-04) to MTYL. Research in SP lab is supported by funding from the NIH (CA258221, and AG068550) and by the Wu Tsai Human Performance Alliance and the Joe and Clara Tsai Foundation.

## Data Availability

The data and code underlying this article are available in the article and in its online supplementary material.

---

**Supp Figure 1.**
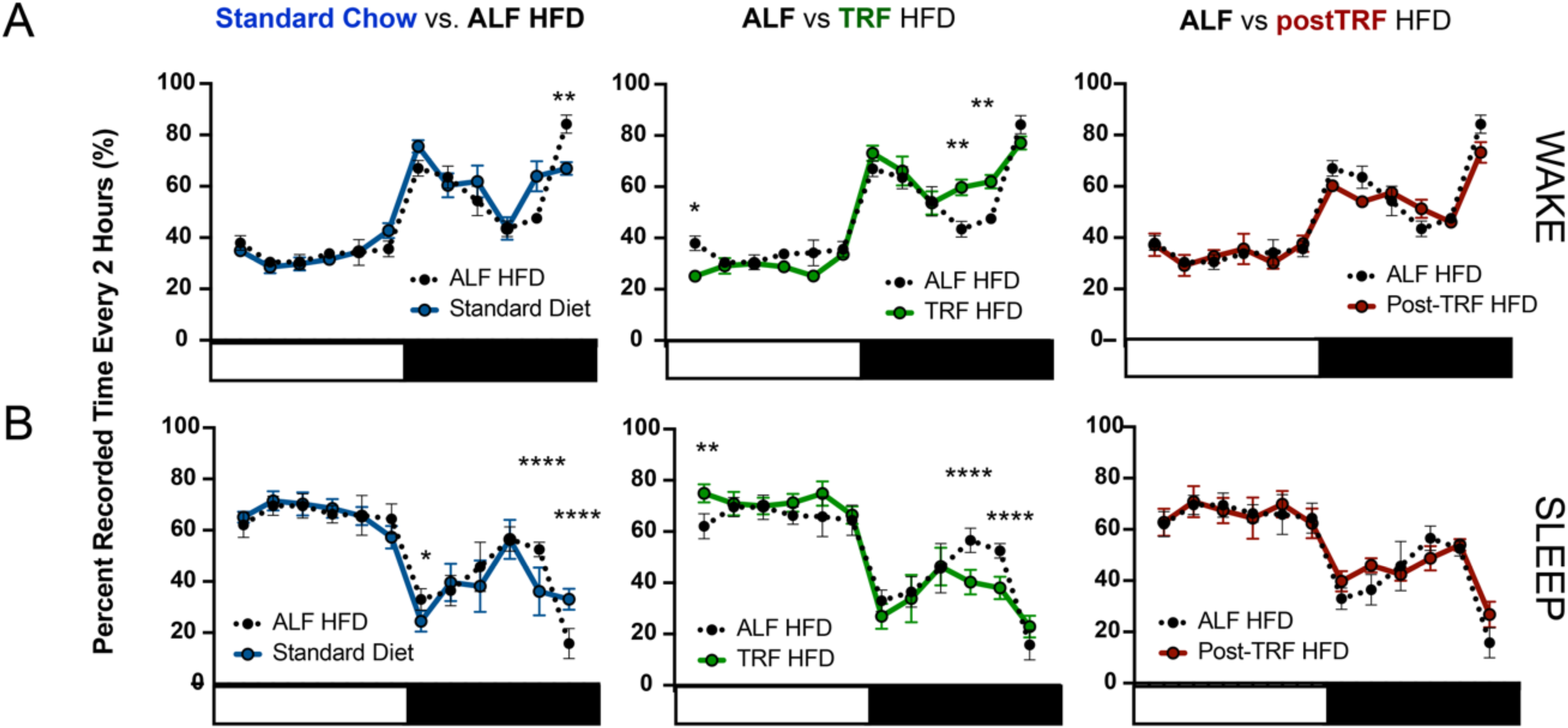
The Effect of Dietary Intervention on Sleep. A-B) Temporal distribution of sleep and wake behavioral states every 2 hours across the 24-hour period. Error bars represent S.E.M. ALF: ad libitum feeding. TRF: time-restricted feeding. HFD: high-fat diet. Adjusted p-value * < 0.05, ** < 0.005, *** < 0.0005, **** < 0.0001.

**Supp. Figure 2.**
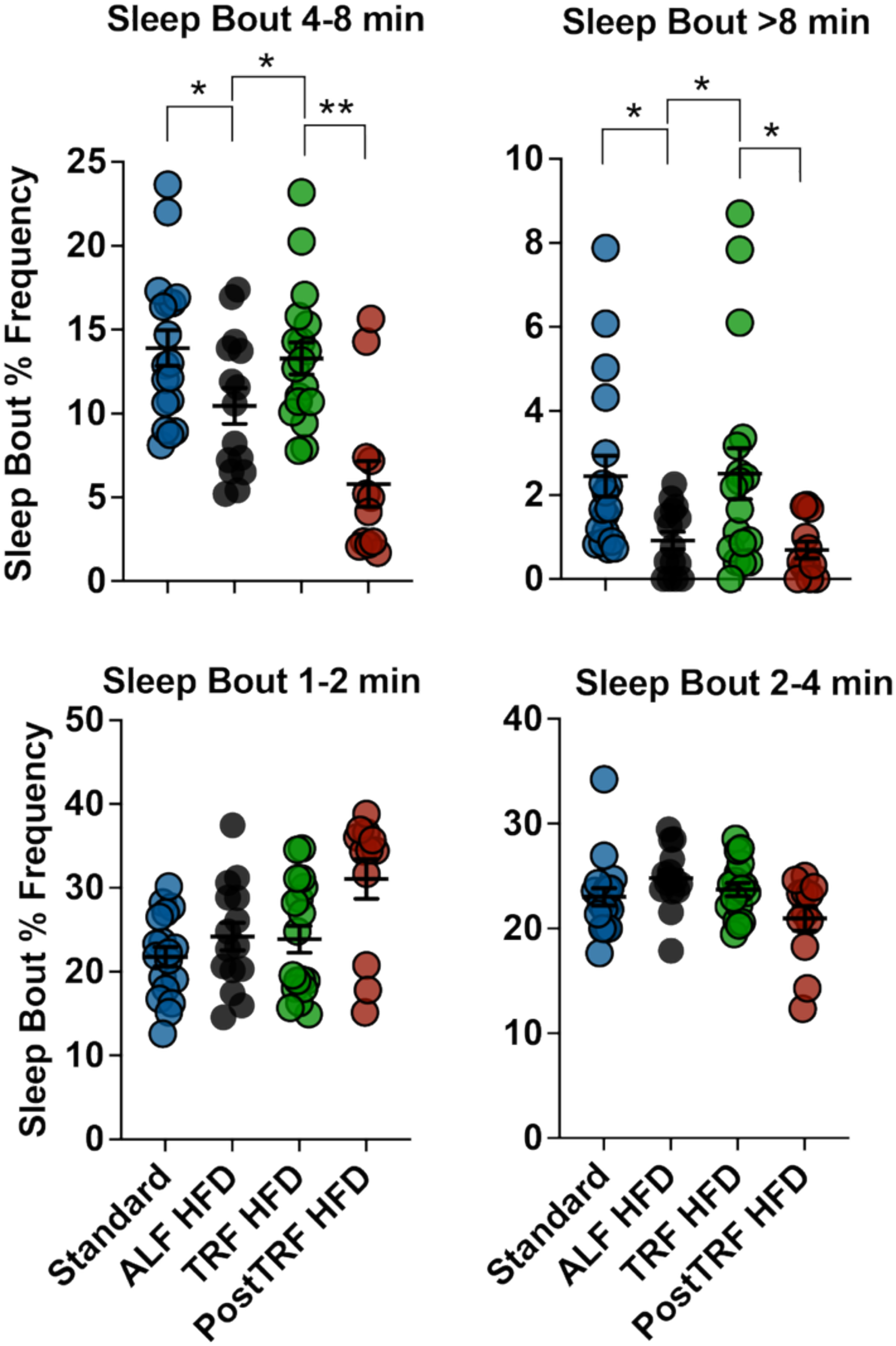
Dietary Effects on Sleep Bout Lengths. The effect of dietary interventions on the percent frequencies of uninterrupted sleep episodes, or sleep bouts, during the light phase (ZT 0 – 12). Changing from a standard chow to a high-fat diet (HFD) decreases the proportion of longer uninterrupted sleep bouts (i.e., 4-8 minutes and >8 minutes). This effect is mitigated by restricting feeding of HFD to the active phase (ZT13-22, TRF HFD). Each dot represents one 24-hour EEG/EMG sleep recording from 4-6 animals. ALF: ad libitum feeding. TRF: Time-restricted feeding. Error bars represent S.E.M. Adjusted p-value * < 0.05, ** < 0.01.

**Supp. Figure 3.**
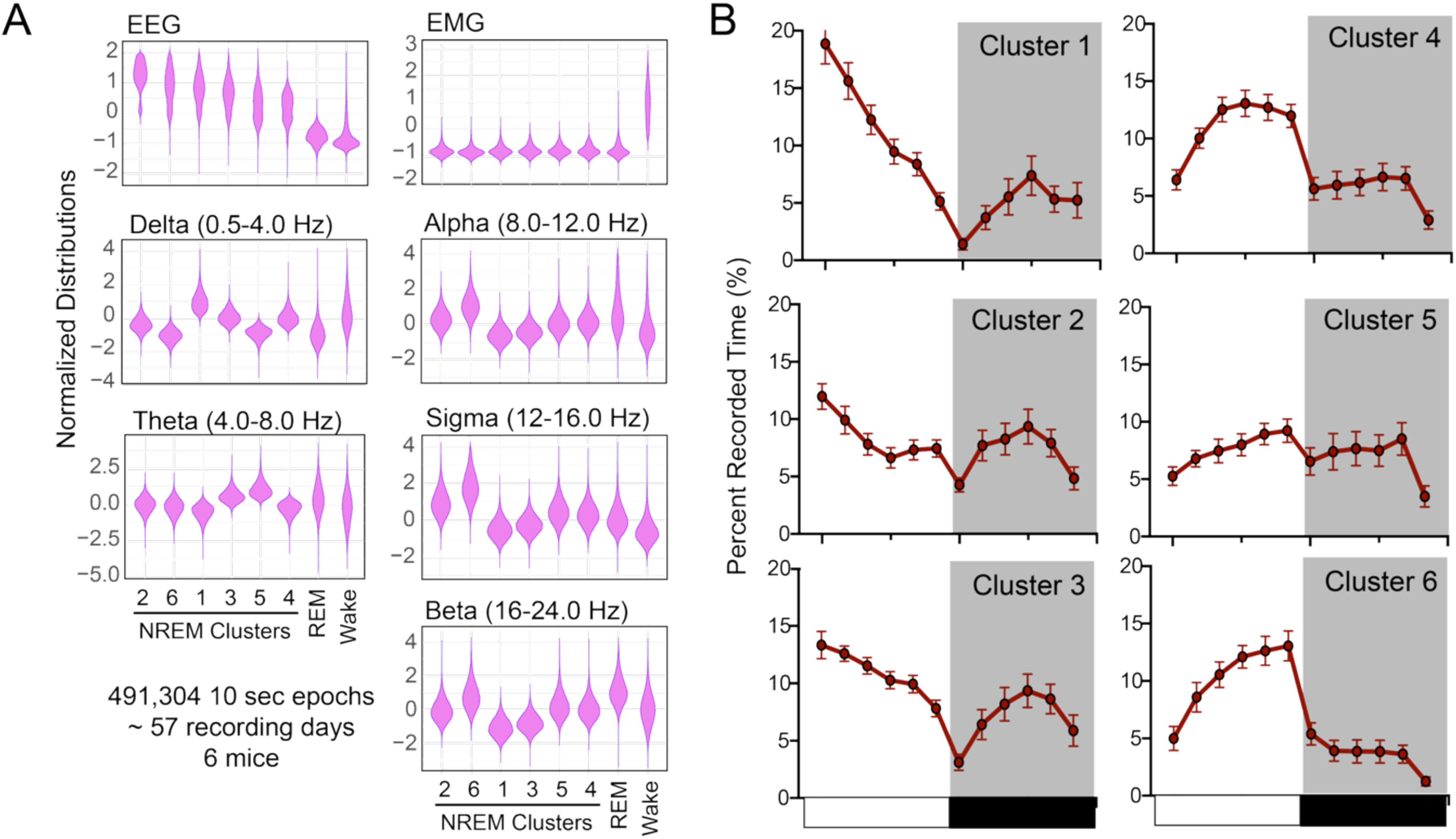
Validation of NREM Substates Feature Characterization in the Dietary Intervention Cohort. The NREM sleep from mice in the dietary intervention cohort was categorized into substates based on electroencephalogram (EEG) and electromyogram (EMG) features defined in the discovery cohort, as presented in the main text (Figure 4). A) Distribution of EEG, EMG, and the power band ratio for Delta, Theta, Alpha, Sigma, and Beta of NREM substate clusters (1-6), REM, and wake. Each of these features was normalized using z-score to the sleep/wake epochs within the 24-hour period. The NREM clusters were ordered on the graph based on the relative descending EEG potential. B) The temporal distribution of NREM Clusters 1-6 across 24 hours. The white bar indicates the light phase (ZT 0-12), and the black bar indicates the dark phase (ZT 12-24). NREM: non-rapid eye movement sleep. REM: rapid eye movement sleep. Error bars represent S.E.M.

